# Designing metabolic division of labor in microbial communities

**DOI:** 10.1101/442376

**Authors:** Meghan Thommes, Taiyao Wang, Qi Zhao, Ioannis Ch. Paschalidis, Daniel Segrè

## Abstract

Microbes face a tradeoff between being metabolically independent and relying on neighboring organisms for the supply of some essential metabolites. This balance of conflicting strategies affects microbial community structure and dynamics, with important implications for microbiome research and synthetic ecology. A “gedanken experiment” to investigate this tradeoff would involve monitoring the rise of mutual dependence as the number of metabolic reactions allowed in an organism is increasingly constrained. The expectation is that below a certain number of reactions, no individual organism would be able to grow in isolation, and cross-feeding partnerships and division of labor would emerge. We implemented this idealized experiment using *in silico* genome-scale models. In particular, we used mixed integer linear programming to identify tradeoff solutions in communities of *Escherichia coli* strains. The strategies we found reveal a large space of nuanced and nonintuitive metabolic division of labor opportunities, including, for example, splitting the TCA cycle into two separate halves. The systematic computation of possible division of labor solutions for 1-, 2-, and 3-strain consortia resulted in a rich and complex landscape. This landscape displays a nonlinear boundary, indicating that the loss of an intracellular reaction is not necessarily compensated by a single imported metabolite. Different regions in this landscape are associated with specific solutions and patterns of exchanged metabolites. Our approach also predicts the existence of regions in this landscape where independent bacteria are viable, but outcompeted by cross-feeding pairs, providing a possible incentive for the rise of division of labor.

## Background

Each microbial cell harbors a finite number of metabolic gene functions. The specific assortment of functions in a given organism is thus the outcome of a tradeoff between the cost of expressing different genes, and the benefit of those genes under different environments. This tradeoff is considered to be one of the possible drivers of diversity in natural microbial communities, giving rise to metabolically differentiated groups rather than an individual superorganism [1–8]. The emergence of metabolically differentiated subpopulations has also been documented to occur from isogenic populations in a fixed environment [9– 19]. The viability of co-existing populations of metabolically differentiated strains or species is often enabled by the exchange of metabolites [7, 11, 15, 16, 20–27]. For example, initially identical populations of *Escherichia coli* evolved on minimal glucose medium have been observed to give rise to a specialized subpopulation of cells that use the acetate secreted as a byproduct of glucose fermentation [15, 20, 21]. More broadly, metabolic interactions mediated by the exchange of small molecules help maintain the diversity and stability of natural microbial communities, and allow communities to accomplish metabolically-intensive tasks [5, 6, 27, 28]. Moreover, obligate metabolic interdependencies (such as mutualism) are believed to contribute to the high prevalence of unculturability and fastidiousness among natural microbial strains [7, 29–33].

A recent and increasingly common strategy to study microbial interdependencies is the construction (or evolution) of artificial microbial consortia specifically designed to display obligate mutualism. Current approaches to building synthetic communities of interacting microbes have so far mainly relied on intuition about simple genetic perturbations that would cause organisms to engage in obligate cross-feeding. In these interactions, one strain is unable to synthesize an essential metabolite (*e.g.*, an amino acid) that is supplied via overproduction or leakage by another strain [34–41]. This ensures that the two strains require each other’s presence in order to grow. While interesting and valuable, these strategies explore only a small portion of the very large and complex space of possible environmental and organismal modifications: in principle, organisms may have the potential to display complex cross-feeding strategies for multiple metabolites simultaneously, or in an environment-dependent manner [42, 43]. In fact, given the complexity of metabolism and its evolutionary history, it is possible that naturally evolved cross-feeding strategies may involve complex metabolic mutualism beyond single amino acid exchanges [44]. In particular, loss of functions in one organism due to compensation by others has been hypothesized to be widespread [45], and may involve multiple genes and complex pathway architectures [25]. In the engineering of consortia for specific metabolic engineering tasks, exploring this larger space of possibilities may open up novel strategies for bio-production.

Surveying the landscape of possible paths for metabolic differentiation leading to obligate mutualism is a combinatorially difficult problem. While future elaborations of existing methods for high throughout genetic modifications (*e.g.*, MAGE [12]) may enable a systematic exploration of this space *in vivo*, computational models can provide a preliminary assessment of the landscape of possible strategies and of how these strategies depend on different constraints on metabolic network complexity. Constraint-based models of metabolic networks, such as Flux Balance Analysis (FBA) [46–56], can specifically be leveraged to ask questions that cannot be easily addressed experimentally. FBA represents metabolism as a set of biochemical reactions, which are inferred from genome annotations and literature curation, and considers cellular metabolism as a resource-allocation problem. Given a set of biochemical, thermodynamic, and environmental constraints, FBA uses linear programming to determine the distribution of fluxes through a reaction network that satisfies a given optimization objective. Typically, this objective is to maximize the flux through a biomass-producing reaction, so FBA determines how a cell should optimally allocate nutrients based on its environment and biochemical capabilities so that growth rate is maximized. FBA has also been increasingly used to study metabolic interactions in microbial consortia [7, 8, 50–54, 57–61, 23–25, 27, 36, 47–49], as well as to predict optimal genetic knockouts for metabolite production [46].

Here, we explore how metabolic differentiation emerges from an isogenic population by using a newly developed constraint-based modeling approach which we name Division Of Labor in Metabolic Networks (DOLMN). In particular, using DOLMN, we explore the space of feasible single-strain or multi-strain metabolic networks by systematically limiting the number of intracellular and transport reactions in each metabolic model. After introducing the mathematical and integer linear programming formulation of DOLMN, we illustrate its capabilities through an analysis of division of labor based on core carbon metabolism in *E. coli* [55]. We next apply DOLMN to a genome-scale *E. coli* model [56], and show that metabolically differentiated and interdependent communities are able to exist under harsher reaction constraints than a single, isolated strain, and even outcompete the single strain in some cases. Our results broaden the scope of possible metabolic interdependencies between metabolically different species, with applications in understanding diversity in natural microbial communities and in designing new artificial consortia.

## Results

### A method to design division of labor in microbial communities

The metabolic division of labor problem consists of partitioning a given metabolic network (which we will refer to as the “global network”) into subnetworks representing individual organisms (which we will also call simply “strains”) (see minimal example in **Figure 1**). Each strain has its own metabolic network, including intracellular reactions, as well as transport reactions, which determine how it interacts with the environment. Environmental availability of different nutrients is defined by constraints on the exchange reactions, which enable the inflow and outflow of environmental metabolites and byproducts. In solving this division of labor problem, we make specific assumptions that reflect the nature and architecture of metabolic networks across different species: (i) We do not set any specific *a priori* expectations about the presence of reactions in different strains. Consequently, a reaction from the global network may be selected to appear in one or more strains, or may not appear in any strain at all; (ii) We expect each strain’s subnetwork to be a well-connected, fully functional metabolism, so as to be capable of producing that strain’s biomass (see Methods); (iii) Upon simulating co-culture of multiple co-existing strains, we require that all such strains must have equal growth rates, so that they would be able to stably co-exist in a chemostat [8, 41, 62, 63].

**Figure 1.**
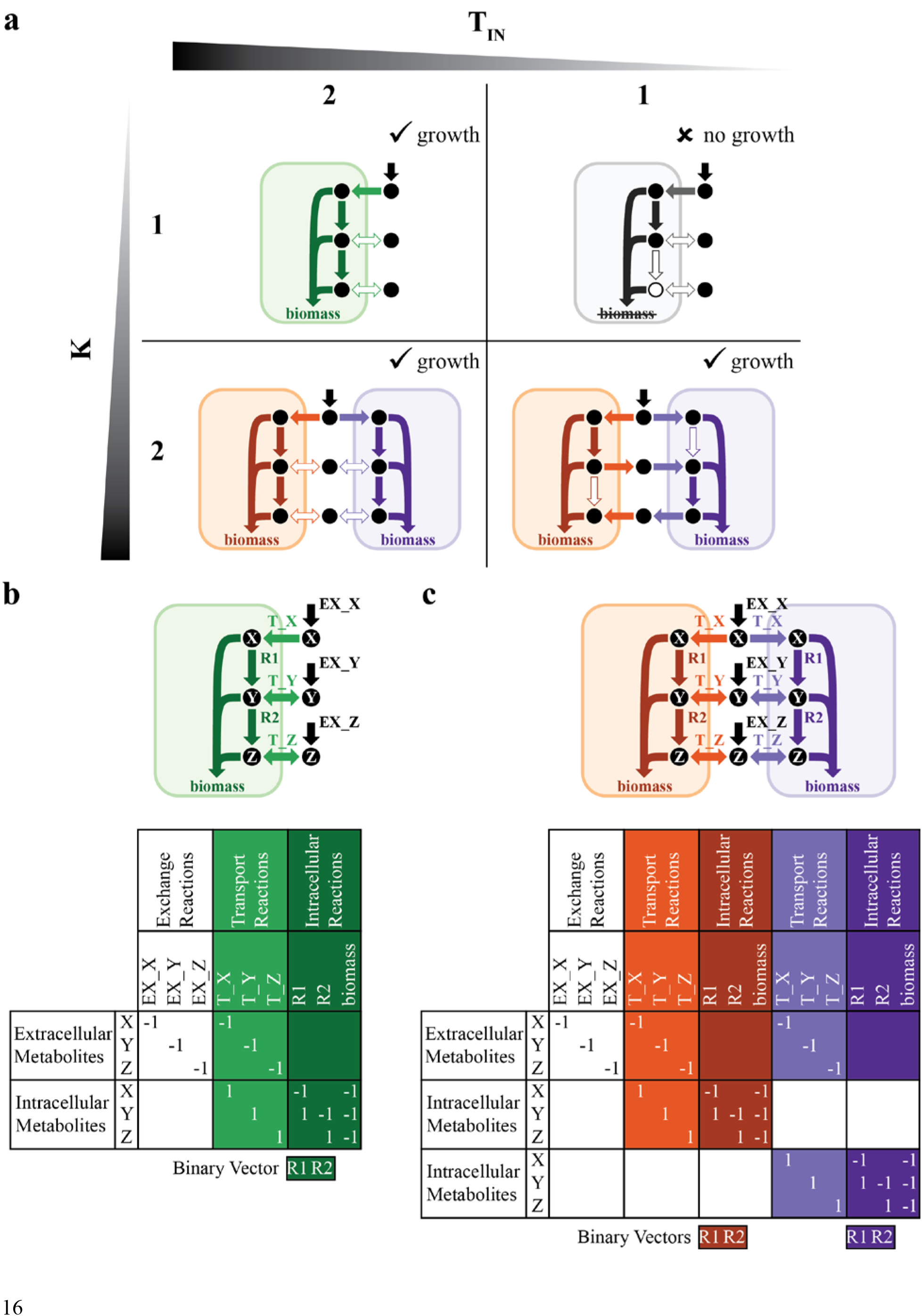
Division Of Labor in Metabolic Networks (DOLMN), illustrated as a toy model. **(a)** One- and two-strain solutions of a toy model. *K* indicates the number of target strains and *T_IN_* is the constraint on the number of intracellular reactions allowed in each strain. Metabolites *X*, *Y*, and *Z* are required for each network to grow (*i.e.*, produce biomass). Two strains can exchange metabolites *Y* and *Z* **(c)**, but a single strain can only take up environmental metabolites **(b)**. **(a)** When *T_IN_*=2, 1- and 2-strain communities perform the same metabolic functions: take up metabolite *X*, convert metabolite *X* to metabolite *Y*, convert metabolite *Y* to metabolite *Z*, and create biomass from metabolites *X*, *Y*, and *Z*. The strains do not take up or secrete metabolites *Y* or *Z*, indicated as a hollow arrow. When *T_IN_*=1, an individual strain is no longer feasible because it cannot create metabolite *Z*, indicated as a hollow circle. The alternative solution, where reaction 1 is knocked out (not shown), is also infeasible because then the single strain cannot create metabolite *Y*. However, 2-strain communities are still feasible because the strains exchange metabolites *Y* and *Z*. If *T_IN_*=1 and the number of transport reactions allowed is constrained to two, then 2-strain communities are no longer feasible (not shown). **(b-c)** Toy metabolic network and corresponding (community) stoichiometric matrices and reaction binary vectors of 1- **(b)** and 2- **(c)** strain communities. The value of each stoichiometric coefficient represents the number of moles of each metabolite that participates in a reaction, with the sign indicating if a metabolite is a product (positive) or a reactant (negative). Exchange reactions are in black; transport reactions are in either light green **(b)**, orange **(c)**, or purple **(c)**; and intracellular reactions are in either dark green **(b)**, orange **(c)**, or purple **(c)**.

To solve this problem, we devised Division Of Labor in Metabolic Networks (DOLMN), which is formulated as a combinatorial optimization problem. Inputs of DOLMN are the global network (encoded in a stoichiometric matrix ***S***, and accompanied by upper and lower flux bounds, as in standard FBA formulations, see Methods); the number of target strains (*K*); and constraints on the number of intracellular (*T_IN_*) and transport (*T_TR_*) reactions allowed in each strain. Key outputs of DOLMN are a binary reaction vector (***t***) whose elements indicate whether a given reaction is present in a given strain, and a continuous flux vector (***x***) for all reaction rates. Note that there is no specific requirement for two or more strains to end up using different reactions from the global network. A specific solution could entail multiple strains having exactly the same reactions, and not interacting with each other. We expect division of labor to arise only upon making *T_IN_* or *T_TR_* too small for any individual strain to be able to survive without receiving specific molecular components from a metabolically distinct partner (**Figure 1a**). Note that elements of ***t*** can switch ON or OFF as a function of the current constraints, irrespective of their state under different constraints.

DOLMN, described in detail in Methods, involves the use of Mixed Integer Linear Programming (MILP). Our problem is NP-complete. It can be solved exactly for core metabolic network models (*i.e.*, global network of ~100 reactions), but it requires heuristics and long computational time for genome-scale models (~1000 reactions).

### Metabolic division of labor in *E. coli* core metabolism

As a first test and illustrative example of DOLMN, we investigated how *E. coli* core carbon metabolism [55] on minimal glucose medium would be partitioned between two strains (*i.e.*, two trimmed versions of the *E. coli* core network) for a given limit on the number of allowed reactions (see **Figure 2** and Methods for details). Besides imposing constraints on the number of reactions allowed in each strain, we further required that both strains have the same growth rate of at least 0.1 hr^-1^, effectively simulating stable co-existence in a chemostat [8, 41, 62, 63]. An interesting outcome of this analysis, obtained for subnetworks containing no more than 11 transport reactions and 24 intracellular reactions, was the discovery of a metabolic strategy in which each strain performs half of the tricarboxylic acid (TCA) cycle. None of the strains, in this case, were able to perform all needed metabolic functions without the inflow of specific metabolites produced by the partner. In particular, exchange of 2-oxoglutarate and pyruvate was necessary for survival of this 2-species consortium (**Figure 2**). This example illustrates how, even for a relatively small network, DOLMN can provide predicted division of labor strategies that could not be easily designed manually. Furthermore, DOLMN could be applied to other core metabolic models [68], which have been generated for a large number of organisms.

**Figure 2.**
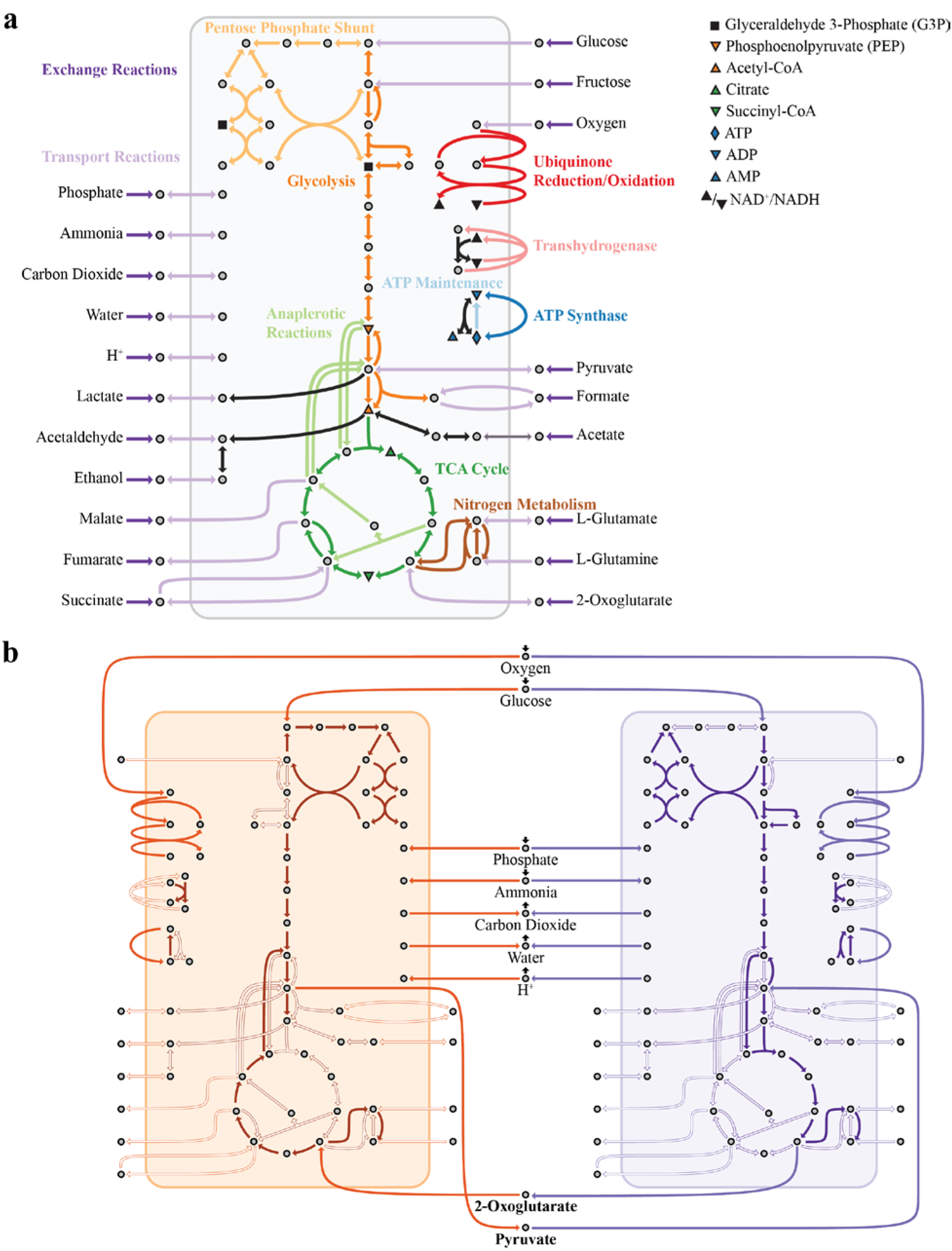
A DOLMN flux solution of E. coli core carbon metabolism. **(a)** The metabolic network of the core *E. coli* model, containing 95 reactions and 72 metabolites. This model contains 20 extracellular metabolites and 52 intracellular metabolites, as well as 20 exchange reactions, 25 transport reactions, 49 intracellular metabolic reactions, and 1 biomass reaction. The biomass reaction is not shown. Pathways and extracellular metabolites are labeled. Key intracellular reactions are labeled using the legend. **(b)** The solution of 2-strain communities when 11 transport reactions (*T_TR_*=11) and 26 intracellular reactions (*T_IN_*=26) are allowed. Reactions that the algorithm identifies as excluded or that have zero flux are indicated as a hollow arrow. The tricarboxylic acid (TCA) cycle is split between the two strains. Both strains consume oxygen, glucose, phosphate, and ammonium, and secrete carbon dioxide and water. The strains exchange the TCA intermediate 2-oxoglutarate and the glycolytic intermediate pyruvate (bolded).

### A growth landscape illustrates division of labor strategies in *E. coli* genome-scale metabolism

We next applied DOLMN to a much larger global network, namely genome-scale *E. coli* metabolism [56]. In this case, individual strains found by the algorithm would represent *E. coli* variants with a reduced set of functionalities. We systematically mapped the landscapes of possible 1-, 2-, and 3- strain simulations to display how the growth rates (**Figure 3a,b** and **Figure S2a** in Additional File 1) vary as a function of *T_IN_* and *T_TR_*. One first observation, consistent with expectations, is that as *T_IN_* decreases (for unconstrained number of transport reactions) individual strains reach a limit beyond which they cannot sustain growth, whereas consortia of two and three strains are still viable. For the example analyzed in **Figure 3**, a 1-strain subnetwork needs at least 254 intracellular reactions to grow, whereas 2-strain subnetworks only require 215 intracellular reactions each, and 3-strain subnetworks require 203 intracellular reactions each (**Figure S1a** in Additional File 1).

**Figure 3.**
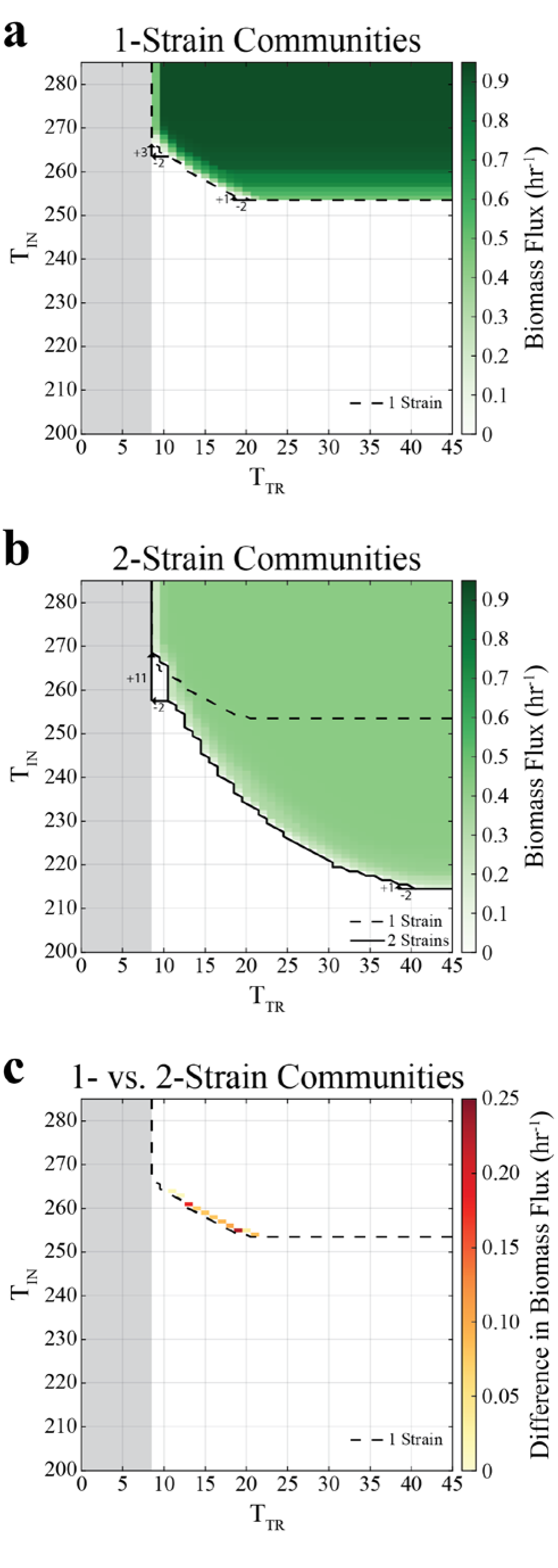
(*T_TR_*,*T_IN_*) growth landscapes of 1- and 2- strain communities. The gray region depicts the infeasible region in which strains cannot grow because there are not enough transport reactions to take up and secrete metabolites. Strains were required to maintain a biomass flux of at least 0.1 hr^-1^. **(a)** Growth landscape of 1-strain communities. Biomass flux values are interpolated to obtain values for when the number of transport reactions allowed (*T_TR_*) are 23, 25–33, 35–37, and 39–45. As less transport constraints are allowed (decreasing *T_TR_*), more intracellular reactions (*T_IN_*) are required. For example, when *T_TR_* decreases from 21 to 19, only 1 additional intracellular reaction is required. However, when *T_TR_* decreases 11 to 9, 3 additional intracellular reactions are required. **(b)** Growth landscape of 2-strain communities. For example, when *T_TR_* decreases from 41 to 39, *T_TR_* only increases by 1, but when *T_TR_* decreases from 11 to 9, *T_TR_* increases by 11. **(c)** Growth landscape of the difference in growth rates between 2- and 1-strain communities, where the constraints at which 2- strain communities grow faster than 1-strain communities are indicated. Only constraints at which 1- and 2-strain communities can both grow are included.

The observed landscapes display a fundamental nonlinear tradeoff between minimizing *T_IN_* (intracellular complexity) and minimizing *T_TR_* (metabolic exchange). This nonlinearity implies that removing the same number of transport reactions at different points along the frontier of the feasible region can be compensated by adding different numbers of intracellular reactions. For example, decreasing *T_TR_* by 2 at large *T_TR_* can be compensated by adding a single intracellular reaction (increasing *T_IN_* by 1), while removing the same number of transport reactions at small *T_TR_* will require a much larger compensation with intracellular reactions (**Figure 3a,b**, **Figure S2** in Additional File 1). It is important to note that decreasing *T_TR_* negatively influences growth because it restricts not only each strain’s ability to take up metabolites, but also its ability to secrete metabolites. If an organism cannot secrete metabolites, it accumulates waste (which results in an infeasible FBA solution). Irrespective of the number of strains in the community, it looks like the *E. coli* strain subnetworks require at least 9 transport reactions in order to support growth (see **Figure S4** in Additional File 1 for which transporters at kept when *T_IN_*=9).

Further analysis of the landscapes for 1-, 2-, and 3-strain communities also reveals the existence of regions in which division of labor potentially provides a competitive advantage. Given that multiple strains co-existing in a consortium have to share available resources, they will tend to grow slower than individually growing strains (**Figure 3a,b**). One notable exception is a thin strip at the boundary in which an individual strain can grow. At this frontier for a single strain, we observe that 2-strain communities can grow more rapidly than 1-strain communities (**Figure 3c**). A biologically important implication of this result is the fact that the 2-strain communities would in principle have the chance to collectively outcompete the 1-strain ones. Similarly, 3-strain communities grow faster than 2-strain communities along the boundary in which 2-strain communities can grow (**Figure S2h** in Additional File 1). These results suggest that the number of strains that achieve the highest growth rates under a given set of circumstances may naturally increase as environmental constraints tighten. This situation could rise, for example, if the burden of protein cost in the cell were to increase, or if selection processes were to gradually favor streamlined strains (*e.g.*, as previously observed experimentally by [10–19, 33]).

### Emergence of obligate mutualism is coupled with sharp metabolic network differentiation and exchange of different chemicals

Based on intuition, and on the results obtained for the core model (**Figure 2b**) we expect that the capacity of two or more *E. coli* strains to survive together under tighter constraints on the number of allowed reactions is due to metabolic division of labor and cross-feeding. A global overview of how metabolism enables co-existence of 2-strain communities in the (*T_TR_,T_IN_*) landscape can be obtained by plotting the metabolic distance (see Methods) between each of the strain’s subnetworks (**Figure 4a**) or the number of metabolites exchanged between them (**Figure 4b**). The fraction of reactions each of the strain subnetworks have in common tends to overall decrease as *T_IN_* and *T_TR_* decrease (**Figure 4a, Figure S5a** in Additional File 1), indicating that it is in general more advantageous for 2-strain communities to perform division of labor as the constraints became more severe. These division of labor strategies are also visible in terms of the number metabolites that the 2-strain communities have to exchange with each other in order to grow (**Figure 4b** and **Figure S6a** in Additional File 1). In short, multiple-strain communities can grow under stricter constraints on *T_IN_* because they distribute metabolic reactions and exchange metabolites. Overall, different 2-strain communities can vary widely in terms of the metabolites being exchanged, with molecules ranging from central carbon compounds such as acetate and pyruvate to amino acids (**Figure 5a**).

**Figure 4.**
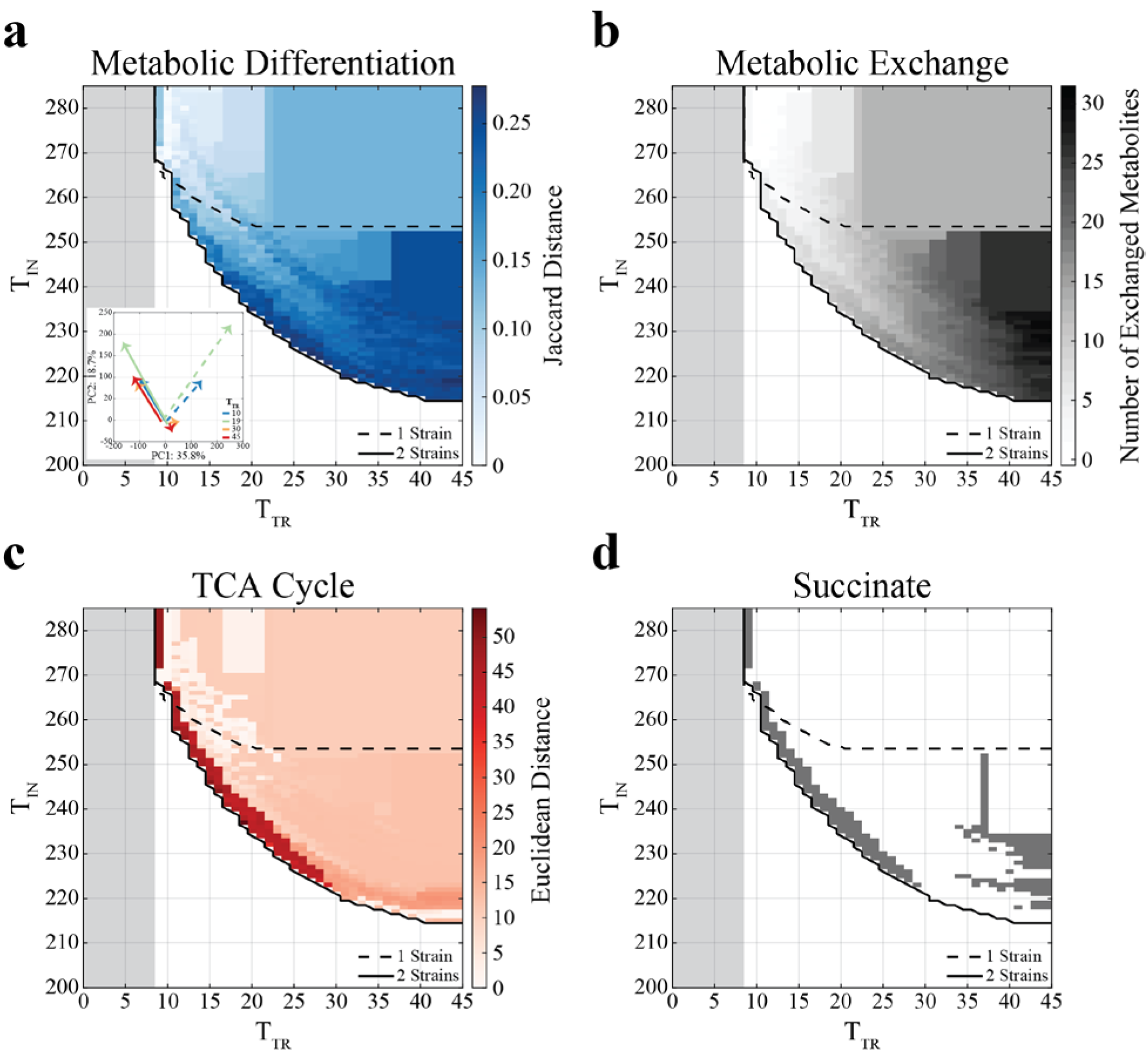
(*T_TR_*,*T_IN_*) landscapes of metabolic differentiation and exchange in 2-strain communities. The gray region depicts the infeasible region in which strains cannot grow because there are not enough transport reactions to take up and secrete metabolites. **(a)** Jaccard distance between reaction binary vectors (***t***) in 2- strain community simulations. Strains that are more metabolically differentiated (have less reactions in common) have a larger distance. Two-strain communities are more metabolically differentiated when they are in the region where a single strain cannot grow. Inset shows the principal component analysis (PCA) plot of 2-strain communities at *T_TR_* = 10, 19, 30, and 45, with arrows pointing from *T_IN_* = 285 to the growth boundary (*T_IN_* = 267, 237, 222, and 215, respectively). **(b)** The number of metabolites exchanged between strains in 2-strain community simulations. The number of exchanged metabolites increases as the constraint on the number of intracellular reactions becomes harsher (decreasing *T_IN_*). As the constraint on the number of transport reactions becomes harsher (decreasing *T_TR_*), the number of exchanged metabolites decreases. **(c)** Euclidean distance between the fluxes of TCA cycle reactions in 2-strain community simulations. **(d)** Constraints at which the 2-strain communities exchange succinate. Two-strain communities distribute the TCA cycle at the same constraints as they exchange succinate (Point-Biserial correlation coefficient of 0.63, **Figure S18a** in Additional File 1).

**Figure 5.**
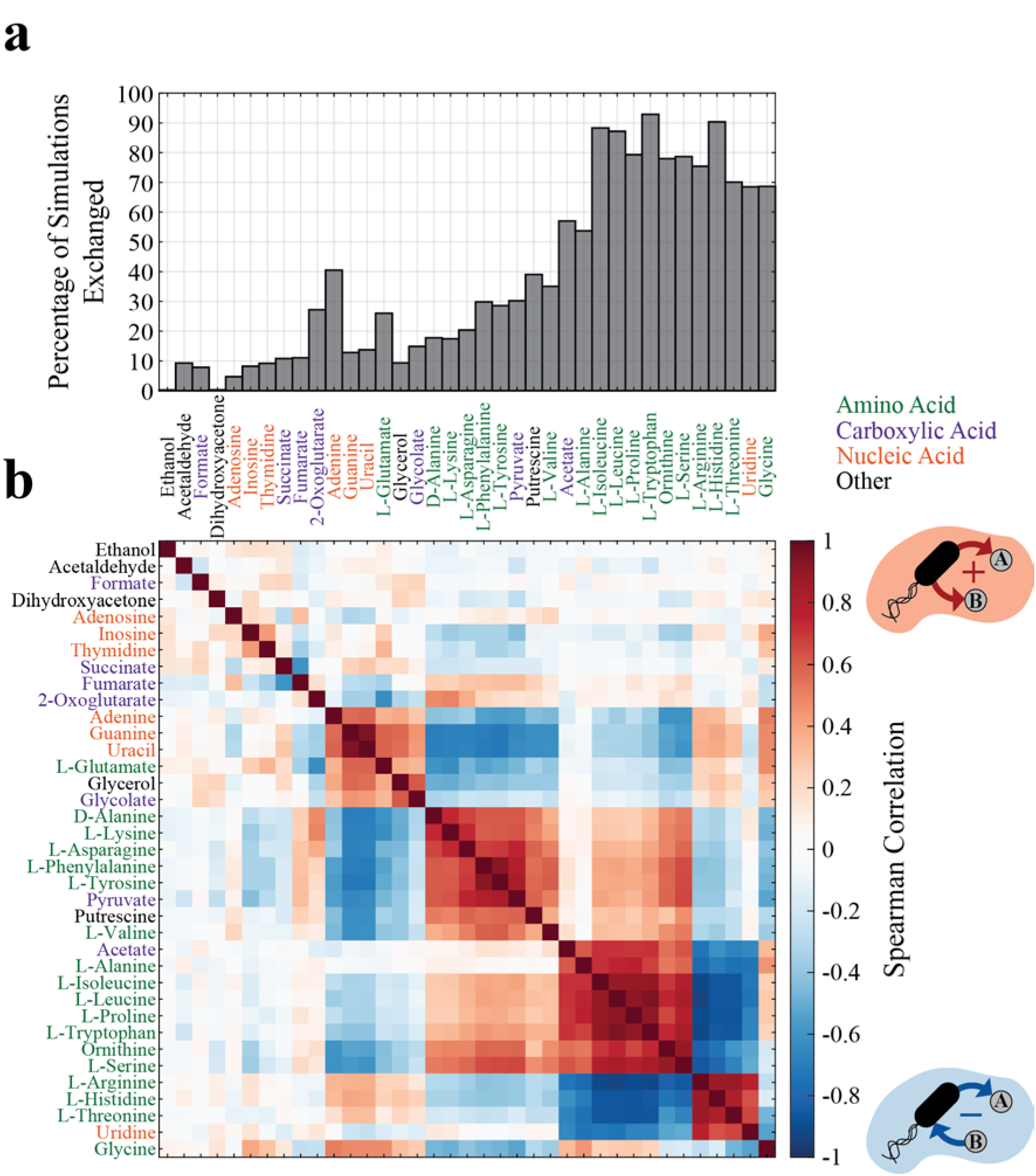
Characterization of exchanged metabolites for 2-strain community simulations. **(a)** Bar plot of the percentage of simulations a metabolite is exchanged in 2-strain communities, clustered hierarchically, as in **(b)**. The Spearman correlation of exchanged metabolites, measuring the relationship of the metabolite exchange. A positive value specifies that the metabolites are both secreted or both taken up by a strain (only secretion is shown) and a negative value specifies that one metabolite is taken up and the other is secreted by a strain.

In order to gain better insight into the metabolic changes that accompany the rise of pairs of obligate mutualistic strains, we reduced the multi-dimensional space of fluxes using PCA (see Methods). Clusters in the principal component space would indicate common metabolic strategies, hardly detectable through visual inspection of the network themselves; paths between these clusters would additionally portray how the strain subnetworks move through this metabolic space as constraints become more stringent. We observed three clusters, indicating three major metabolic strategies (**Figure 4a inset**, **Figure S8** in Additional File 1). The largest cluster occurs at the origin, and corresponds to large *T_IN_*. That is, for fairly unconstrained intracellular metabolism, all strains (for 1- and 2-strain simulations) perform the same metabolic strategy, in line with the expected metabolic regime for *E. coli* grown on minimal glucose medium [55, 56]. However, as *T_IN_* becomes more constrained (*i.e.*, the number of intracellular reactions allowed decreases), the 2-strain subnetworks move away from each other, indicating that they diversify into different metabolic strategies. Note that the specific path followed by these networks as *T_IN_* decreases is also a function of *T_TR_* (**Figure 4a inset**, **Figure S8** in Additional File 1).

To understand the different metabolic strategies associated with different clusters in the PCA, we calculated the Euclidean distance between pathways in each of the two strains in a 2-strain community and mapped it to the (*T_T_*_R_,*T_IN_*) landscape (see Methods). A striking outcome of this distance analysis was the fact that, for a broad range of *T_TR_* values (9–29), the 2-strain subnetworks had a large difference in their TCA cycle (**Figure 4c**), mirroring the observation reported for the core *E. coli* network (**Figure 2b**). This strategy of splitting the TCA cycle was also supplemented by large differences in glycolysis/gluconeogenesis, which was in fact observed along the entire *T_IN_* boundary for 2-strain communities (**Figure S9** and **Figure S10** in Additional File 1), suggesting that this could represent a generalized version of the acetate utilization phenotype observed in classical evolutionary experiments [8, 20, 21].

As indicated by the increase in number of exchanged metabolites as constraints become tighter (**Figure 4b**), the pairs of metabolically differentiated strains described above can survive due to metabolic cross-feeding. For example, in a 2-strain community, one of the strains could utilize the metabolites available from the environment, and secrete byproducts that can enable the other strain to survive. We again used the (*T_TR_,T_IN_*) landscape to track how constraints affect the metabolites being exchanged (**Figure 4d, Figure S12** in Additional File 1). Different metabolites display drastically different patterns (**Figure S12** in Additional File 1): some metabolites are exchanged almost universally (*e.g.*, acetate) while others appear only in specific sub-regions of the landscape (*e.g.*, glutamate below the *T_IN_* boundary for 1-strain communities). Notably, succinate is shown to be exchanged predominantly at the *T_IN_* boundary of 2-strain communities (**Figure 4d**), in very close correspondence to the area of the landscape where the TCA cycle is drastically split (**Figure 4c**). This suggests that, based on genome-scale simulations, succinate would be one of the key intermediates for the rise of *E. coli* strains surviving by using complementary halves of the TCA cycle.

### Fluxes of metabolic sharing can be strongly coupled

Consortia of obligate symbiotic partners are predicted to emerge in regions of the (*T_TR_*,*T_IN_*) landscape where 1-strain communities are infeasible. As shown above, what makes these consortia viable (*i.e.*, what makes it feasible for the corresponding strains to produce biomass despite the strong restriction on intracellular reactions) is the possibility of metabolite exchange between the 2-strain communities.

We thus sought to perform analysis of the metabolites being exchanged across the whole (*T_TR_*,*T_IN_*) landscape. In the specific setup used for the *in silico* experiments, out of 143 total extracellular metabolites, only 37 metabolites (25.9%) were exchanged in at least one simulation of 2-strain communities. Most of these exchanged metabolites (78.4%) were exchanged in at least 10% of all co-culture simulations (**Figure 5a**). A class of abundantly exchanged molecules is the set of amino acids. These solutions can be viewed as similar to artificially imposed auxotrophies used to engineer synthetic consortia [37–41, 64]. Other frequently exchanged molecules include carboxylic acids (*e.g.*, acetate and pyruvate) and nucleic acids (*e.g.*, thymidine and adenine), which are known to be exchanged in natural communities [20, 21, 27, 59].

Additional insight can be gathered by exploring the relationship between exchanged metabolites, *i.e.*, by asking whether we should expect specific pairs of metabolites to be simultaneously exchanged in the same or opposite directions between two organisms. Knowledge of such correlations/anti-correlations may be useful as a strategy for choosing biomarkers (if two organisms exchange A, they are also likely to exchange B), as an indicator of fundamental metabolic tradeoffs (X can provide A only if Y provides B), or as a broad suggestion for the existence of unavoidable couplings in the interactions present in a microbial community. We applied the Spearman correlation to exchange reaction fluxes (**Figure 5b**) in order to measure if metabolites are exchanged jointly (both taken up or secreted by a strain, positive *ρ*) or reciprocally (one is taken up and the others is secreted by a strain, negative *ρ*). The Spearman correlation can tell us if metabolic complementation exists between 2-strain communities.

Two major patterns emerge from this analysis. First, by looking at the hierarchically clustered correlation matrix, one can immediately recognize several block structures. The largest block structure seems dominated by amino acid exchange. In particular, two sets of (mostly) amino acids seem to be highly correlated within each set, but highly anti-correlated across the sets. The first set includes L-arginine, L-histidine, L-threonine, and uridine, which are all correlated with each other, whereas the second set includes L-isoleucine, L-leucine, ornithine, L-proline, L-serine, L-tryptophan (as well as acetate and L-alanine, with weaker coefficients). Interestingly, these anti-correlated block sets do not seem to map trivially to different precursor pathways (*e.g.*, upper vs. lower glycolysis), suggesting that other metabolic tradeoffs may determine these patterns. One can also observe a second block structure dominated by amino acids that are all correlated, containing D-alanine, L-asparagine, L-lysine, L-phenylalanine, pyruvate, L-tyrosine, and – to a less extent – fumarate, 2-oxoglutarate, putrescine, and L-valine.

The second significant outcome of this correlation matrix is related to the TCA cycle splitting result illustrated above. In particular, succinate and fumarate are anti-correlated (*ρ* = -0.60), indicating reciprocal exchange. These two metabolites are intermediates of the TCA cycle, and are involved in sequential steps, suggesting that this anti-correlation corresponds to the metabolic division of labor strategy characterized by splitting of the TCA cycle (**Figure 2b**). This is further conformed by the fact that the region in the (*T_TR_*,*T_IN_*) landscape where succinate is exchanged matches very closely with the region in which the 2-strain communities have very distinct use of the TCA cycle (**Figure 4c,d**).

## Discussion

In this study, we proposed a set of methods and thought experiments to systematically explore the space of possible division of labor strategies in synthetic microbial consortia as the number of transport and intracellular reactions are constrained. We found not only that 2-strain communities can survive with less metabolic reactions than individual strains because they distribute reactions and exchange metabolites, but also that under some conditions, two cross-feeding strains may grow faster than one strain. There is a nonlinearity in how loss of internal reactions can be compensated by transporters, indicating that a single metabolite supplied by the environment or another strain can offset the loss of multiple intracellular reactions. These non-trivial synergistic states have not been hypothesized before and would be difficult to find without the aid of computational methods.

Although this study is purely computational, it provides a new conceptual framework and new predictions, including specific testable modifications, and it enables us to perform analyses that are currently beyond current experimental capabilities. The phenotypic space we explore involves 5,000 *in silico* experiments on the full *E. coli* network, across 1-, 2-, and 3-strain communities at multiple constraints on the number of transport and intracellular reactions. Furthermore, for each organism’s model we take into account a large number of variables (1,075 reactions and 761 metabolites in each strain). Efficient algorithms are therefore a key step towards designing functional and stable division of labor strategies in synthetic microbial communities.

Experimental testing of the proposed division of labor strategies in *E. coli* may prove challenging due to the multiplicity of gene deletions that would have to be simultaneously performed. Recently developed technologies, such as multiplex automated genome engineering (MAGE) [65], conjugative assembly genome engineering (CAGE) [66], and the CRISPR/Cas system [67], could in principle facilitate such an endeavor. Still, a challenge of implementing multiple targeted knock-outs experimentally is the chance of encountering high-order epistatic interactions between genes that are difficult to predict computationally, and that may result in non-viable strains. Thus, instead of implementing all of the genetic perturbations at once, it may be advisable to engineer increasingly complex interactions involving gradual modifications of different reactions and pathways. Moreover, rather than deleting the genes, one could consider engineering promoters to reduce the flux through each reaction and potentially let the strains evolve in the lab.

Our results may be relevant for microbial ecology of natural communities, as well as for the study of synthetic microbial consortia. Ongoing efforts in synthetic ecology are currently mostly focused on engineering metabolic dependencies by making one strain unable to produce a terminal biomass precursor (*e.g.*, an amino acid), which is then provided by another strain. Although this strategy has yielded new insight into how microbes interact within communities, it may not be reflecting the possible complexity of natural metabolic interactions. Obligate mutualistic organisms within communities may be thermodynamically and metabolically linked based on the exchange of metabolites that are part of core metabolic processes. Our approach has the capacity to uncover this kind of “deep symbiosis” between organisms, such as the split TCA cycle. Interestingly, the TCA cycle appears in an incomplete form in many microbes [68–70], and reduction in the lower half of the TCA cycle occurred in *E. coli* evolution experiments [71, 72]. Moreover, some bacteria, such as *E. coli*, switch between the full TCA cycle and a branched variant when operating under aerobic and anaerobic conditions, respectively [73, 74]. These observations also suggest that complex – and yet poorly understood – solutions to community-level metabolic efficiency may have arisen through co-evolution in the form of intricate division of labor strategies.

Our predictions were initially produced under the assumption that the different perturbations applied by our method correspond to genetic modifications. However, our predictions could equally be interpreted as being a consequence of instances of gene down-regulation instead of gene loss [14]. In other words, all the solutions found by DOLMN may in principle manifest themselves in the form of phenotypic differentiation within a population of cells. In a complex multicellular system (such as the human body) this could mean division of labor among cell types in different tissues, whereas in clonal populations of microbes this could imply phenotypic variation due to heterogeneous gene expression [75–77]. Single-cell studies of microbial physiology [14], aided by genome-scale models of metabolism, could help unravel both genomic and transcriptional variability potentially associated with division of labor in the microbial world.

## Conclusions

We computationally explored the possible ways in which sets of metabolic reactions can be distributed among interacting microbial strains, with the goal of better understanding the tradeoff between metabolic self-reliance and mutualistic exchange. The mathematical problem of designing metabolically viable organisms and communities from a global set of possible reactions is a very difficult one. The approach, heuristics, and examples illustrated in this work show, however, that solutions identified by our algorithm are feasible and biologically interpretable. In addition to illustrating specific nontrivial avenues for engineering communities of co-dependent *E. coli* strains, our approach could be viewed as a first step towards addressing a broader, overarching question, namely whether the metabolic network of individual organisms in natural communities is predictable based on first principles. One could imagine, in particular, that abundant horizontal gene transfer and long-term selection processes in ecosystems may have acted over geological time to efficiently allocate genes into mutually dependent organisms, very much like our algorithm does. With increasing computational power and further optimized algorithms, it may become possible to extend our approach to a larger number of organisms, with the potential of providing a general theoretical scaffold for understanding how environments shape division of labor strategies and microbial diversity.

## Methods

### Flux balance analysis (FBA)

To mathematically formulate FBA, let **S** denote the stoichiometric matrix of dimensions *m*×*n*, where *m* is the number of metabolites and *n* the number of metabolic fluxes. Metabolic fluxes are defined as a vector ***x***, where ***x_lb_*** and ***x_ub_*** are lower and upper bounds, respectively, on the metabolic fluxes. These bounds are implied by empirical evidence of irreversibility or by nutrient availability in the growth medium. The cellular objective is expressed as a vector of weight coefficients for each reaction (*e.g.*, biomass), denoted by **c**, and the optimal objective value is a scalar *Z_opt_*. The FBA problem is formulated as:

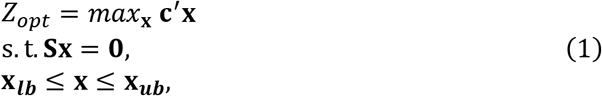

where **0** is the vector of all zeroes and primes indicate transpose.

### Community-level flux balance analysis

In order to perform FBA capturing all reactions spanning an entire microbial community, we introduce a “universal stoichiometric matrix,” also denoted by **S**, which expresses the stoichiometric coefficients of all metabolic reactions in the community irrespective of the organism they belong to, as in [61]. Specifically, **S** ∈ ℝ^*M×N*^ where *M*=*M_e_*+*M_i_* represents the number of distinct metabolites and *N*=*N_e_*+*N_t_*+*N_i_* represents the number of distinct reactions (see **Figure S16** in Additional File 1). The *M* distinct metabolites consist of two types: *M_e_* extracellular and *M_i_* intracellular metabolites. There are 3 different types of reactions: *N_e_* extracellular reactions, *N_t_* transport reactions, and *N_i_* intracellular reactions. The availability of nutrients (extracellular metabolites) from the environment is encoded in the extracellular reactions, and intracellular reactions encode each organism’s metabolism. Organisms use transport reactions to move metabolites between their intracellular compartment, which is unique to each organism in the community, and the extracellular environment, which is shared by all organisms in the community.

### A method for metabolic division of labor

We first reformulate the universal stoichiometric matrix **S** to construct putative stoichiometric matrices for each species in the community. In particular, we construct a community stoichiometric matrix **S^c^** whose structure is shown in **Figure S17** in Additional File 1. The block matrices **S^e^**, **S^t1^**, **S^t2^**, and **S^i^** in **S^c^** are consistent with those in **S**. Organisms in the community share the same nutrients and extracellular reactions. Because there are *K* organisms in the community, we replicate the block [**S^t2^,S^i^]** that includes transport reactions and intracellular reactions *K* times and diagonally arrange them in **S^c^**. Similar compositions of stoichiometric matrices had used in previous work on community level flux balance modeling [43, 78, 79].

After the intracellular metabolites are obtained via the transport reactions, intracellular reactions take place inside each organism. This construction leads to a community stoichiometric matrix 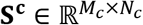, where *M_c_*=*M_e_*+*K*×*M_i_* and *N_c_*=*N_e_*+*K*×(*N_t_*+*N_i_*) Notice that **S^c^** has one block column for extracellular reactions (*N_e_* columns) and *K* block columns (of dimension *N_t_*+*N_i_*), one for each organism, including all transport and intracellular reactions.

To capture design choices, we introduce a binary putative vector ***t*** = (*t*_1_, … *t_Nc_*), where *t_j_* ∈ {0,1}, *j*= 1, …, *N_c_*, is a binary variable, indicating whether the *j*-th reaction is included in the corresponding organism (**Figure S17** in Additional File 1). With ***t*** and **S**^c^ available, we can partition **S**^c^ to *K* individual matrices, **S**^k^, by removing column *j* with *t_j_*= 0.

The problem of identifying ***t*** can now be formulated as the following MILP problem:

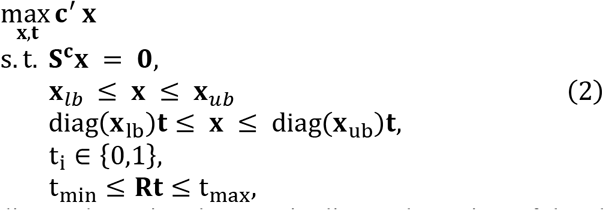

where diag(**x**) denotes a diagonal matrix whose main diagonal consists of the elements of vector **x**, **R** is a regularization matrix, and **t**_min_, **t**_max_ are appropriately defined constant vectors. Specifically, by appropriately defining **R**, **t**_min_, **t**_max_, we can impose constraints on the number of internal and transport reactions for each organism.

### *E. coli* models

*E. coli* core [55] and genome-scale iJR904 [56] models were retrieved from the BiGG database [80]. The models were downloaded as .mat files in the COBRA (COnstraints-Based Reconstruction and Analysis) format [81]. Stoichiometric matrices were reformatted as a community stoichiometric matrix **S**^c^, as previously described and shown in **Figure S17** in Additional File 1. Reaction and metabolite names were re-ordered to correspond with the community stoichiometric matrix.

### The First Optimization Problem

Suppose there are *K* organisms. The upper bound on the number of active transport and intracellular reactions in each organism is T_TR_ and T_IN_, respectively. We let x_biomk_ denote the flux of the biomass reaction for each organism *k*= 1, …, *K*. We also let *TR_k_* and *IN*_k_ denote the index sets of the transport and internal reactions for each organism *k*, respectively. The MILP (**Problem 2**) takes the form:

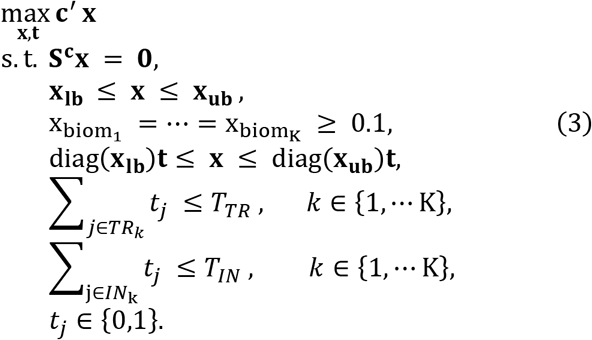

Let **x**, **t^*^** denote an optimal solution of the MILP problem above.

We note that solving such a large-scale MILP, which involves hundreds or thousands of integer variables, is computationally expensive. Our experiments suggest that solving Problem 3 for a community of two *E. coli* core models can be done relatively quickly (on the order of minutes or hours). On the other hand, solving the problem for a community model of two iJR904 *E. coli* models is very time consuming. We employ certain methods to speed up finding an optimal solution. One method leverages the fact that instances with similar values for T_TR_ and T_IN_ have similar sets of active reactions. Specifically, as we decrease the values of T_TR_ and T_IN_ we use the sets of active reactions corresponding to larger T_TR_ and T_IN_ to generate feasible solutions that are offered as putative solutions to the MILP solver. This tends to drastically decrease solution times. A complementary approach uses a decomposition idea. In particular, for solving problems involving a community of *K* organisms, we can use solutions for *K*-1 organisms and append solutions for the additional organism. This generates feasible solutions and it is possible to search for an effective feasible solution by varying the way the *K*-organism community is decomposed into a (*K*- 1)-organism community and an additional organism.

### The Second Optimization Problem

In order to reduce redundant fluxes in transport and intracellular reactions, a second optimization problem is introduced where the integer variables are fixed to the optimal solution **t**\*of the first stage problem (**Problem 3**), and the biomass fluxes are also set to the optimal values obtained by the first stage problem (**Problem 3**). Specifically, we solve:

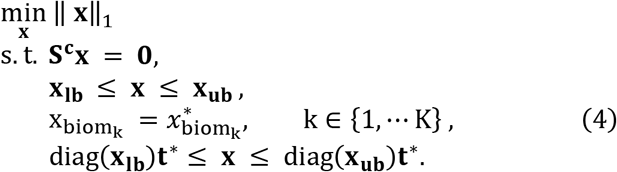

This problem minimizes the ℓ1 norm of the flux vector to induce sparsity (as in the “sparse FBA” of [81]) and can be rewritten as a linear programming problem.

### Data Structure and Analysis

For each constraint on the number of transport reactions, *T_TR_*, DOLMN outputs a structure, 𝓒 (“model”) for each strain. This structure 𝓒 contains fields to indicate the constraint on the number of intracellular reactions, *T_IN_* (“model.sparse_con”); the reaction names in the global network (“model.rxns”); the growth rates at each *T_IN_* (“model.biomass”); the reaction flux values at each *T_IN_* (“model.flux”); and the reaction binary integer values at each *T_IN_* (“model.int”). The function algorithm2models parses 𝓒 into individual metabolic models, calculates the exchange flux of each individual strain (instead of the community exchange flux), and identifies exchanged metabolites.

The analysis of the DOLMN output is performed in several MATLAB scripts. All analysis is split between the core and full iJR904 *E. coli* model. The scripts dolmn1a_parse_iJR904.m and dolmn1b_parse_core.m apply the function *algorithm2models* to all of the DOLMN outputs for the full and core iJR904 *E. coli* models, respectively. The scripts dolmn2a_summary_iJR904.m and dolmn2b_summary_core.m restructure the data for plotting, calculate the Jaccard distance and exchange flux correlations, and perform standard PCA (MATLAB function *pca*). Interpolation for the constraint landscapes are performed in dolmn2a_summaryInterp_iJR904.m and dolmn2b_summaryInterp_core.m. All main text figures are plotted by dolmn3_figures_main.m and all supplementary figures are plotted in dolmn4_figures_supp.m.

All data (raw and analyzed), functions, and scripts can be found in the GitHub repository (https://github.com/segrelab/dolmn). Analysis was performed with MATLAB 2017b. MILPs were solved by GUROBI 7.0 (http://www.gurobi.com, [82]).

### Transport Reaction Flux to Exchange Reaction Flux

Metabolite exchange requires (i) that the metabolite was not already in the environment (was not part of the medium); (ii) that one organism secretes the metabolite (positive exchange flux); and (iii) that another organism takes up the metabolite (negative exchange flux). DOLMN finds the net exchange flux of the community, so we had to calculate the exchange flux of each strain in a multi-strain solution. The calculated exchange flux is the sum of all transport reactions:

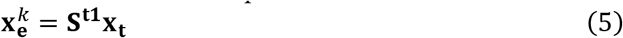

and the sum of each calculated exchange flux must be equal to the net exchange flux of the community:

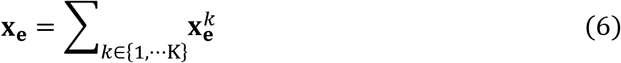

### Calculating Metabolic Differentiation

The Jaccard distance was used to measure the difference between models based on how many reactions were different between strains. The Euclidean distance was used to measure how the flux distributions differed between strains. The built-in MATLAB function *pdist2* was used to calculate Jaccard and Euclidean distances, with the ‘jaccard’ and ‘euclidean’ metrics, respectively.

### Identifying Exchanged Metabolites

Metabolites are exchanged if one strain secretes the metabolite (has a positive exchange flux) and the other takes it up (has a negative exchange flux). Metabolites that were part of the medium (*e.g.*, hydrogen ions) were not considered to be exchanged even if one strain secretes it and the other takes it up. We created the MATLAB function *identifyExchangedMets* to determine which metabolites, if any, are exchanged between strains.

### Principal Component Analysis

Principal component analysis (PCA) was performed on the intracellular flux values of 1- and 2- strains using the built-in MATLAB function *pca*. The biomass flux was excluded from the intracellular reactions and was instead used to normalize the intracellular flux values.

### Spearman Correlation of Exchange Reaction Flux

The Spearman correlation of exchange reaction fluxes and the phi coefficient of exchanged metabolite profiles were calculated using the MATLAB function *corr*, with the ‘Spearman’ and ‘Pearson’ type, respectively. While the Spearman correlation yields information on how metabolites are exchanged together (*e.g.*, if they are both secreted or taken up or if they are inversely exchanged), the phi coefficient indicates if a metabolite is exchanged at the same (*T_TR_*,*T_IN_*) constraints. In general, the phi coefficient measures the association between two binary variables (in this case, the metabolite exchange profiles as in **Figure 4d** and **Figures S11-12** in Additional File 1). The Spearman correlation of exchange reaction fluxes was clustered by the shortest Euclidean distance using the MATLAB functions *linkage* and *dendrogram*.

## List of Abbreviations Used

FBA: Flux Balance Analysis
DOLMN: Division Of Labor in Metabolic Networks
LP: Linear Programming
MILP: Mixed Integer Linear Programming

## Declarations

### Ethics approval and consent to participate

Not applicable.

### Consent for publication

Not applicable.

### Availability of data and material

All data generated and analyzed in this study and the corresponding code are available in the GitHub repository (https://github.com/segrelab/dolmn).

### Authors’ contributions

All authors contributed to the design of the study and to the definition of the problem. TW, QZ, and IP formulated and solved the optimization problem. TW and IP developed the heuristics for large datasets. TW generated all simulation datasets. MT and TW analyzed the data. MT and DS implemented the metabolic analyses and biological interpretation. MT, TW, IP, and DS wrote the manuscript. All authors read and approved the final manuscript.

### Acknowledgements

We are grateful to Sara Collins and Joshua Goldford for discussions on division of labor in microbial communities, and to all members of the Segrè Lab for helpful discussions and feedback.

### Description of additional data files

The following additional data are available with the online version of this paper:
**Additional File 1:** Supplemental Figures and Tables

